# Blue light stimulated-autofluorescence green flash of lysosome-related organelles in the intestinal cells of nematodes

**DOI:** 10.1101/2023.10.16.562538

**Authors:** Chieh-Hsiang Tan, Keke Ding, Mark G. Zhang, Paul W. Sternberg

**Affiliations:** Division of Biology and Biological Engineering, California Institute of Technology, Pasadena, CA 91125, USA; Innoland biosciences, Hangzhou, 310000, China

**Keywords:** lysosome-related organelles, gut granules, autofluorescence, light stimulation, Caenorhabditis elegans, Steinernema hermaphroditum, Oscheius tipulae, Panagrellus redivivus, Pristionchus pacificus

## Abstract

The lysosome-related organelles (“gut granules”) in the intestinal cells of many nematodes, including *Caenorhabditis elegan*s, play an important role in metabolic and signaling processes, but they have not been fully characterized. We report here a previously undescribed phenomenon in which the autofluorescence of these granules displays a “flash” phenomenon in which fluorescence decreases are preceded by sharp increases in fluorescence intensity that expand into the surrounding area when the granules are stimulated with blue light. Autofluorescent granules are present in the intestinal cells of all six nematode species examined, with differences in morphology and distribution pattern. Five species exhibit the flash phenomenon: *Panagrellus redivivus* (Clade IV), *Steinernema hermaphroditum* (Clade IV), *C. elegans* (Clade V), *Oscheius tipulae* (Clade V), and *Pristionchus pacificus* (Clade V). The reaction of the granules to blue light stimulation greatly differs among different developmental stages and may also be dependent on physiological conditions. In addition, even within the same animal, the sensitivity of individual granules differs, with some of the variation associated with other characteristics of the granules, such as their overall location within the intestine. We hypothesize that the differences in response to blue light indicate the existence of different sub-populations of gut granules in nematode intestines, and the visually spectacular dynamic fluorescence phenomenon we describe might provide a handle on their eventual characterization.

## Introduction

The intestinal cells of *Caenorhabditis elegans* (Maupas 1900) contain a type of organelle known as gut granules, suggested to be the rhabditin granules observed in many other nematodes (Maupas 1900; Cobb 1914; Hedrick 1935; Chitwood and Chitwood 1950; Thomas and Quastler 1950; Laufer et al. 1980). The birefringent and autofluorescent gut granules are robustly present in intestinal cells and intestinal precursor cells (Laufer et al. 1980; Hermann et al. 2005) and are so prominent that they have served as useful markers for the intestinal cell lineage in *C. elegans* development. Based on morphological, biochemical, and genetic evidence, these granules are considered lysosome-related organelles (LROs; Clokey and Jacobson 1986; Kostich et al. 2000; Hermann et al. 2005; Delevoye et al. 2019). While far from being fully characterized, previous studies found these organelles to be associated with metabolism and homeostasis processes such as the storage of fat, cholesterol, and heme (Schroeder et al. 2007; Lee et al. 2015; Sun et al. 2022) and the storage and sequestering of trace metals (Roh et al. 2012; Chun et al. 2017). Gut granules are also known to be involved in the biogenesis of small signaling molecules, such as ascarosides, and in the resistance to stress and pathogens (Le et al. 2020; Hajdu et al. 2023).

In this study, we found that some autofluorescent granules in nematode intestines underwent a rapid and dynamic change in fluorescence intensity when observed in the GFP channel (blue light excitation; green emission). In *C. elegans*, this phenomenon depends on the worm’s ortholog of human Rab32/38, *glo-1* (<u>G</u>ut granule <u>LO</u>ss) (Hermann et al. 2005; Morris et al. 2018), which has been shown to be required for gut granule biogenesis, suggesting that gut granules were the source of the autofluorescent green flash. High-resolution confocal imaging of *ex-vivo*, partially exposed *C. elegans* intestines revealed the spatial-temporal patterns of the fluorescence intensity changes, which we found to be different under the green and red channels. We reasoned that this suggests the existence of at least two distinct types of fluorophores or fluorophore transitions in the gut granules. Green fluorescence intensities in the granules sharply increased, rapidly diffused, and then dissipated, a phenomenon that we describe as “gut granule green flash.” No similar rapid increase in red fluorescence intensities was observed prior to their simultaneous dissipation. By increasing the intensity of the (GFP) excitation wavelength blue light during imaging, we were able to significantly reduce the amount of time between the start of the light exposure and the onset of gut granule green flash, suggesting that the phenomenon is likely induced by blue light. We found that their responses to blue light stimulations differ greatly among different developmental stages, in which we were able to reliably document the phenomenon in the second (L2), fourth (L4), and the stress-resistant dauer larvae stages, but were unable to do so in the L1 stage and adult animals. The association of the phenomenon with the availability of food in *ex-vivo* settings suggests that it may also be influenced by physiological conditions. To further understand the green flash phenomenon, we investigate the autofluorescent granules in five other nematode species. Including two other clade V nematodes (based on the five major clade classification of Blaxter et al. (1998)), namely *Oscheius tipulae* (Lam and Webster 1971) and *Pristionchus pacificus* (Sommer et al. 1996), as well as three clade IV worms, namely *Steinernema hermaphroditum* (Stock et al. 2004), *Panagrellus redivivus* (Linnaeus), and a species of the *Panagrolaimus* genus (*Panagrolaimus sp.* PS1159) (Sommer and Sternberg 1996). We found autofluorescent granules to be present in the intestinal cells of all five species, four of which exhibit the green flash phenomenon with different patterns. From these observations, we conclude that it is likely that, at least in some developmental stages, gut granules comprise distinct subgroups, literally highlighted by fluorescence dynamics.

## Results

### Autofluorescent lysosome-related organelles in the intestine of *C. elegans* displayed dynamic changes in fluorescent intensities when exposed to blue light

When observed with the GFP channel, the autofluorescent granules in the intestine of *C. elegans* can undergo dynamic changes in fluorescence intensity (Figure 1; Video S1, S2). Specifically, there is a sharp increase in fluorescence intensity in the granules, followed by a quick dissipation and diffusion into the surrounding area (Figure 1). This “flash” behavior of the autofluorescent granules can be observed in some but not all developmental stages. We observed the events in the stress-resistant dauer stages larvae (15 of 20 worms) (Figure 1 A-E; Video S1), the second stage (L2) larvae (5 of 5 worms), and the fourth stage (L4) larvae (19 of 20 worms) (Figure 1F-H; Video S2), but not in the first stage (L1) larvae (0 of at least 5 worms) and adult animals (0 of 5 worms).

**Figure 1.**
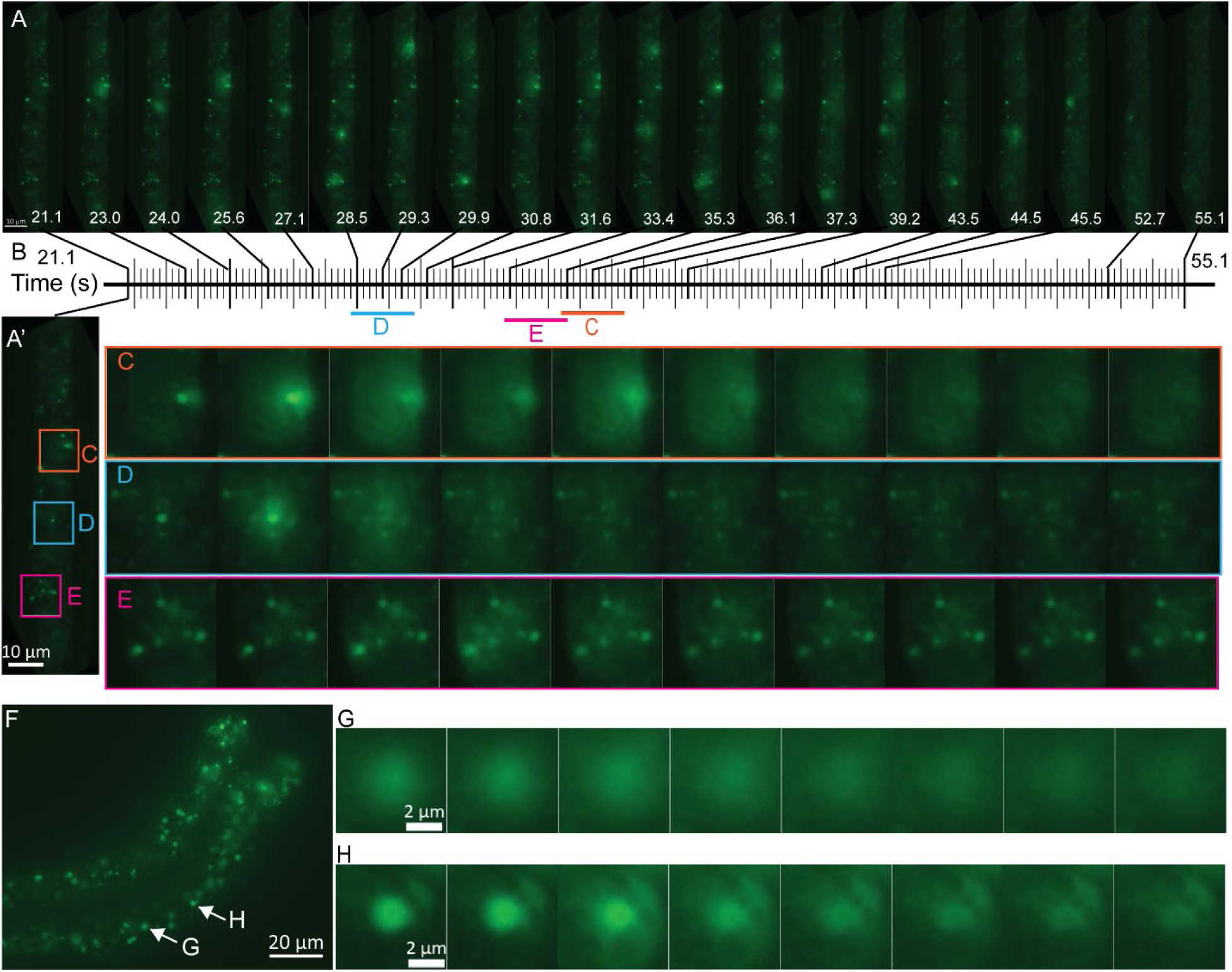
Autofluorescent granules in intestinal cells displayed dynamic changes in fluorescence intensities. (A-E) Dynamic fluorescence intensities change in intestinal granules of a dauer stage larva. (A) Fluorescence images (eGFP channel) of the intestine of a dauer stage larva of wild-type (N2) *C. elegans* at different time points during the analysis. Values represented are seconds and start (from 0) with the initiation of live imaging. The complete series of images is presented in Video S1. (A’) An enlarged image of A at time 0, with sections represented in C-E indicated with boxes. Due to the movement of the animal, the positions of the granules were not exactly the same as in C-E. (B) A timeline positioning the images of A to E. Each vertical line represents an individual time point with an image taken. The images were acquired at a rate of ∼5 frames per second. (C-E) Enlargement of sections of the intestine shown in A, focusing on individual granules at the time of rapid intensity changes over periods of ∼1.8 seconds. In each case, the dissipation of the granules’ autofluorescence was preceded by an increase in intensity (2nd panel in C and D, 3rd in E) and then a “diffusing” cloud that increased the fluorescent intensities of the surrounding areas. In (C), fluorescence intensity changes in the 5th panel may be due to a second granule undergoing the process in a different focal plane. In (E), the granule can still be clearly observed after the event, albeit with decreased fluorescence intensities. (F-H) Fluorescent dynamics in intestinal granules of an L4 stage larva. (F) Fluorescent images (eGFP channel) of the intestine of an L4 stage larva of wild-type (N2) *C. elegans*. (G, H) Enlargement of sections of the intestine shown in F, focusing on individual granules at the time of the dynamic changes over periods of about 1.1 seconds. The complete series of images can be found in Video S2. Scale bar: (A, A’): 10µm; (F): 20µm; (G, H): 2µm.

We reasoned that these autofluorescent granules are lysosome-related organelles known as gut granules (Laufer et al. 1980). We examined *glo-1(lf)* worms deficient in gut granule biogenesis and observed neither these granules nor the flash phenomenon in the *glo-1(lf)* worms (L4, 0 of 11 worms) (Hermann et al. 2005), suggesting that the granules with the flash phenomenon are gut granules. We therefore refer to the phenomenon as “gut granule green flash”.

We hypothesized that the phenomenon was induced by the blue light used for excitation since it only occurs after the start of the imaging process (Video S1, 2). Indeed, increasing the intensity of the blue light significantly shortens the amount of time it takes for the flash phenomenon to occur in the intestine of wild-type dauer stage larvae-45.4 seconds (Range: 41 – 54, n = 5), when the intensity was set at 10% Maximum); but down to 15.2 seconds (Range: 13 – 19, n = 5) and 12 seconds (Range: 9 – 14, n = 5), when the intensity was set at 50% and 100%, respectively.

### At least two types of fluorophores are likely present in the gut granules with distinct fluorescence dynamics

To investigate the dynamics of gut granule green flash, we extruded the intestines of L4 stage hermaphrodites by cutting the animal open posteriorly near the anus (Figure 2G) and imaged the intestine using scanning confocal microscopy. Under this *ex-vivo* experimental setting, we were able to observe and characterize the fluorescence dynamics with increased resolution. However, in contrast to the *in-vivo* setting, we found gut granule green flash to be rare under this condition. Part of the difference could be due to the intensities of the stimulating light. Since the primary function of the intestine is food digestion, we reason that autofluorescent granules’ fluorescence flashes could be associated with nutrient uptake. In *C. elegans*, food availability significantly influences the rhythmic defecation cycle (Thomas 1990), controlled by calcium oscillations in the intestinal cells (Dal Santo et al. 1999). To test whether gut granule green flash is associated with food availability, we provided food (*Escherichia coli* OP50) to the experimental animal on microscopic slides (Figure 2G). We found that in worms with a significant amount of food near the head region, the occurrence of the gut granule green flash increased dramatically (Figure 2A-D; Video S3). All 7 recordings of worms provided with food displayed the phenomenon (Figure 2C, D, I), while only 1 of 7 without food did, and at a low frequency (Figure 2A, B, I). It is unclear why the autofluorescent granules’ fluorescence flash phenomenon in the *ex-vivo* setting was strongly correlated with food availability, but it may imply that gut granule green flash could be a part of or a consequence of a metabolic or signaling process in food intake. A possible explanation for the difference in food dependency in the *in-vivo* and *ex-vivo* settings is that only in the *ex-vivo* setting, with no provision of food, were the intestinal cells removed from the usual content of the intestinal lumen. In subsequent experiments in which we examined gut granules’ fluorescence flash in *ex-vivo* intestines, we provided food to increase the frequency of the flash.

**Figure 2.**
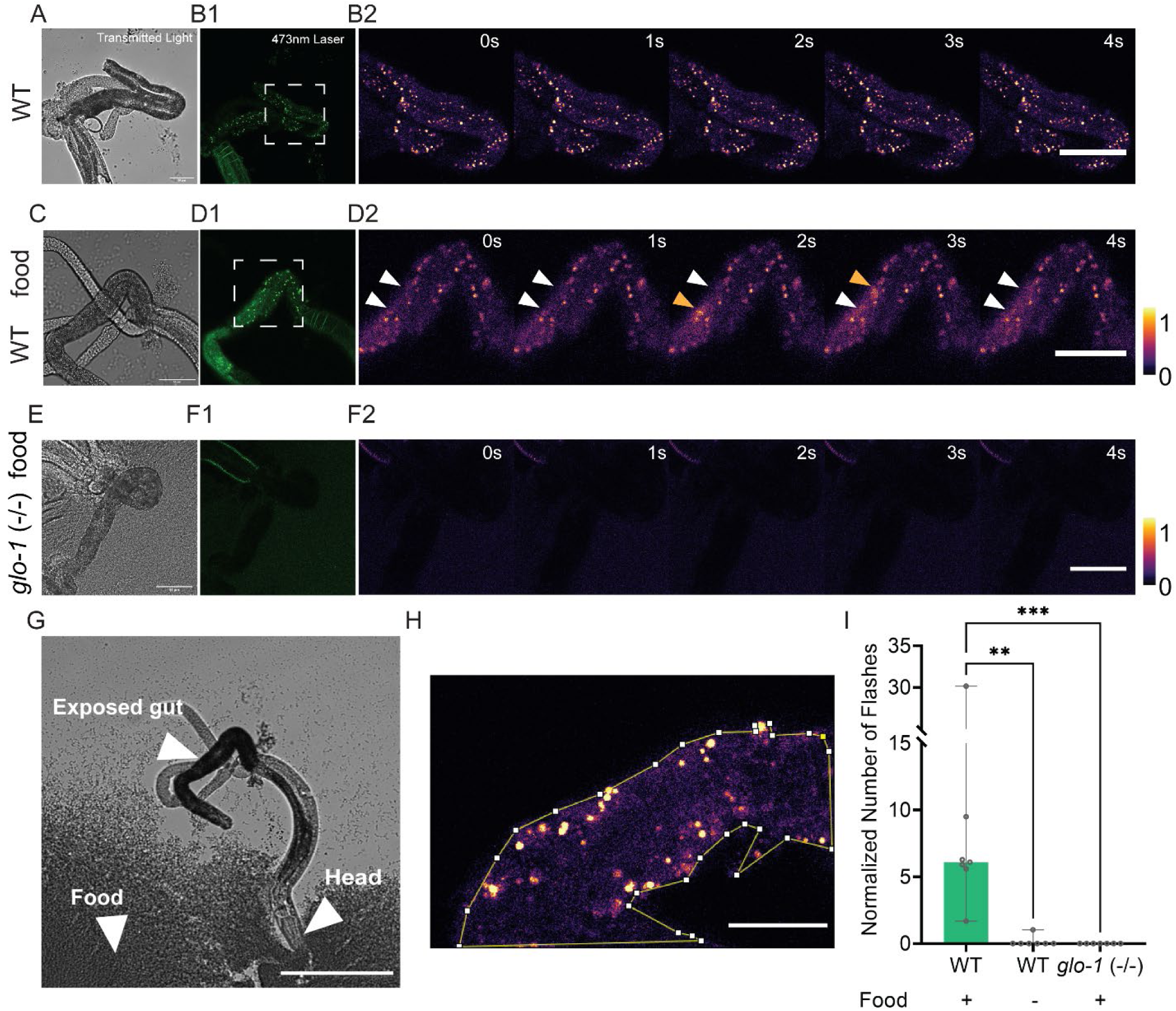
Gut granule green flash is correlated with food availability in the *ex-vivo* experimental setting. The field of view was illuminated and recorded with transmitted light (A, C, E), epifluorescence (B1, D1, F1), and 473nm laser illumination (Ex: 473nm; Em: 500-550) (B2, D2, F2). In (B2, D2, F2), 5-second clips of time series with 473nm laser illumination were pseudo-colored and shown. (A-B) In wild-type worms not provided with food on the slide, the phenomenon was rarely observed. (C-D) In wild-type worms provided with food on the slide, the occurrences of the phenomenon increased dramatically. Orange arrowheads in (D2) indicate the positions of gut granule green flash events. White arrowheads indicate the positions of those events before and after the flash. (E-F) No gut granule fluorescence and no events were detected in *glo-1(lf)* worms, even in the presence of food. (G) An example of the *ex-vivo* experimental preparation. L4 stage nematodes were carefully incised near the anus to expose the gut. In the “with food” conditions, nematodes were positioned to embed their heads in food. (H-I) Gut granule green flash events were manually identified and analyzed. To normalize the number of events for comparison, the counts for each sample were temporally averaged in the area of exposed intestine within the field of view, as shown in (H) (details in Materials and Methods). (I) Normalized data from all conditions were plotted (N=6-7), and each group was compared with the wild-type (WT) + food group (**P<0.01, ***P<0.001, Mann-Whitney U test). The vertical bar indicates the data range. Scale bar: (A-F): 50 µm; (G): 200 µm; (H): 20 µm.

To further test whether these autofluorescent granules that flash are gut granules, we analyzed the *ex-vivo* intestines of *glo-1(*lf) mutant animals. *glo-1(*lf) animals displayed a large reduction in the number of autofluorescent granules in the intestine, similar to what was described by Hermann et al. (2005), and also completely eliminated detectable flashes (Figure 2E, F, I). None of the seven *glo-1(*lf) animals displayed food-induced flashes of fluorescence, as seen in all seven wild-type animals (Figure 2I), suggesting that the autofluorescent granules that flash are gut granules.

When wild-type worms under the *ex-vivo* setting were provided with food, we observed that the green fluorescence, illuminated with the 473nm laser (Ex: 473nm; Em: 500-550) of the confocal microscope, rose sharply and significantly (Figure 3A-C; Video S3). The intensity increase is not only limited to the original “core” area (the granule proper, as identified by the high level of fluorescence prior to the event) but also to the surrounding areas (which we refer to as the “cloud;” details of how the area is identified are in Methods; Figure 3D). The fluorescence intensity in both the “core” and the “cloud” subsequently dropped off, with the intensity at the “core” decreasing to a level lower than baseline, and the intensity in the “clouds” recovered to the baseline (Figure 3D). Nevertheless, when illuminated with the 561nm laser (Ex: 561nm; Em: 580-680), the emitted red fluorescence lacked the initial sharp increase observed in green fluorescence but fell concurrently (Figure 3E). We estimated that the rapid changes in fluorescence intensity occur within 10 seconds. The differences observed in the dynamics of green and red fluorescence intensities suggest that they likely represent two distinct types of fluorophores. It is also possible, however, that they represent a single fluorophore with two distinct transitions.

**Figure 3.**
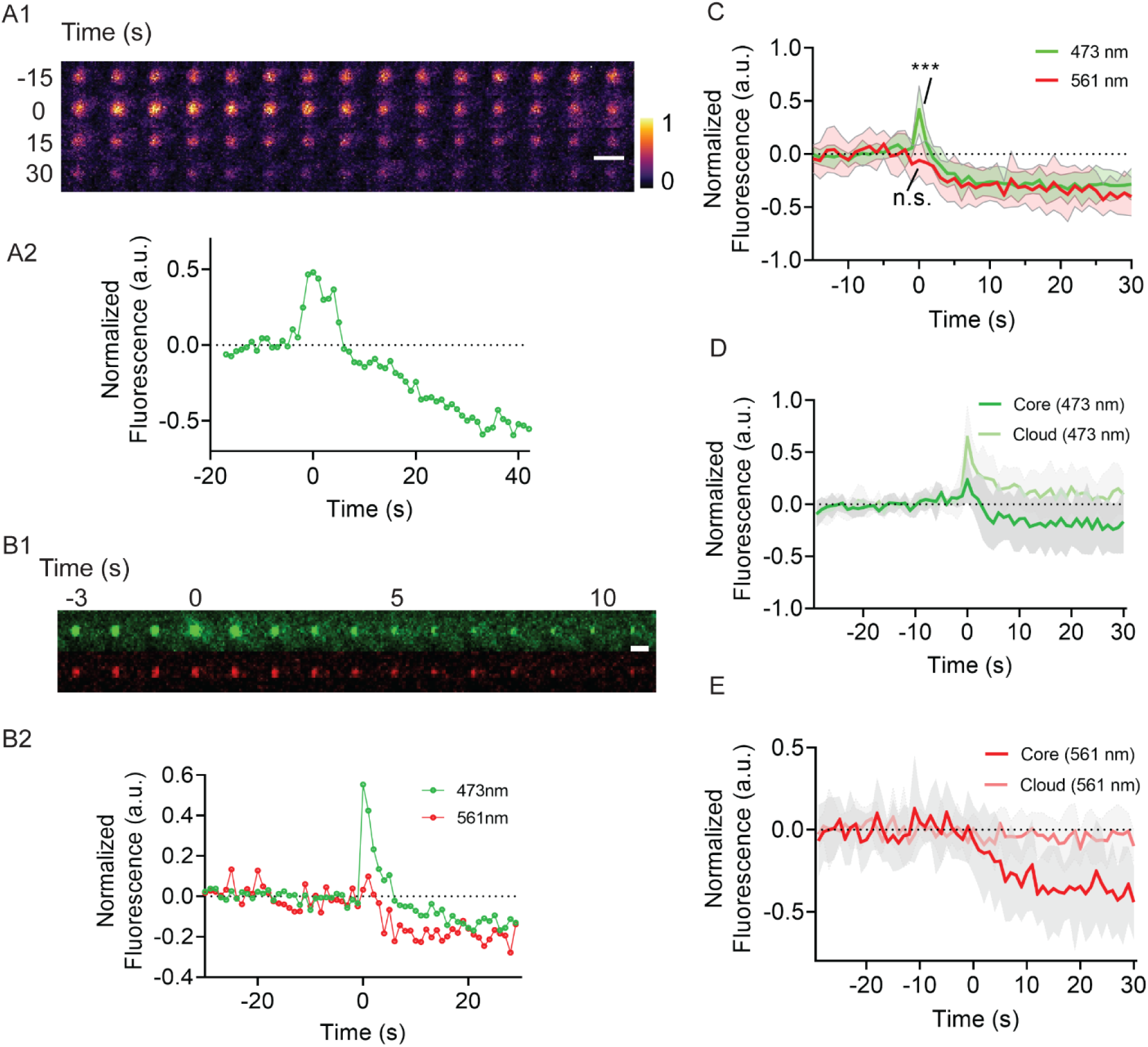
The dynamics of gut granule green flash. (A) The green fluorescence in the granule increases and falls sharply. Representative example of changes in green fluorescence during gut granule green flash. (A1) A 1-minute pseudo-color time series of a gut granule green flash event. Imaging rate: 1Hz. The time axis is zeroed at the onset of changes in fluorescence (see Methods). (A2) Normalized green fluorescence intensity plot of the same granule shown in (A1). (B-C) The green fluorescence in the granule increases and falls sharply, while the red lacks the initial sharp increase. (B) A representative example of two-color time-lapse imaging (B1) and normalized quantifications (B2). In A2 and B2, the fluorescence intensities of each granule are normalized by the baseline fluorescence calculated as the mean of fluorescence in the ROIs prior to the changes in fluorescence. (C) The average fluorescence intensity of all recorded green flash events in one sample. The green fluorescence shows a distinct peak after time 0, while the red fluorescence does not. (D-E) Gut granule flashes. Two-color imaging of all recorded gut granule green flash events in a representative sample. The normalized fluorescence of each data point at time 0 in the green channel is compared to baseline (Normalized Fluorescence=0) and shows a statistically significant difference (***), while a parallel comparison between red channel data point vs. baseline does not show a significant difference (n.s.). (D-E) The granular area (“core”), as well as the surrounding ring area that presents diffusion of fluorescence (“cloud”), were measured separately in both 473nm (Ex: 473nm; Em: 500-550) (D) and 561nm (Ex: 561nm; Em: 580-680) (E) illumination. (D). The intensity increase in green fluorescence is not only limited to the original “core” area but also to the surrounding “cloud.” (E) No intensity increase was observed with the red fluorescence, neither in the “core” nor the “cloud.” Raw images for analysis were identical to (C), except a longer pre-onset timeframe was selected for normalization. In (C-E), pooled values are presented as mean ± s.d (N=11 in D, N=8 in D and E). The time axis is zeroed at the onset of changes in fluorescence. Scale bar: 5 µm.

One possible mechanism behind the green flash phenomenon is the breakdown of gut granule membranes induced by the energy carried by the blue light from the GFP channel. We, therefore, explore whether exposing the worm under the shorter wavelength DAPI channel would give a similar result. We found that continuous exposure to DAPI channel light turned the intestine/worm blue (Video S4), but a similar fluorescence flash phenomenon was not observed (tested in the intestine of wild-type dauer stage larvae, with 10%, 50%, and 100% intensity as with the previous experiment with GFP channel; at least five animals were tested in each setting). One of the fluorophores residing in the gut granules has been identified as anthranilic acid (Babu 1974; Coburn et al. 2013), which has been associated with the increase in the blue fluorescence near the death of the animals (“death fluorescence”). Although the fluorescence changes described in this study are distinct both spatially and temporally from the death fluorescence reported, it is still possible that the breakdown of gut granule membranes causes both.

In spite of its lack of an obvious light-sensing organ, *C. elegans* exhibits negative phototaxis (Ward et al. 2008; Edwards et al. 2008; Bhatla and Horvitz 2015). Given our finding that gut granules may be light sensitive, we wondered whether their presence is associated with the animal’s light response. We found that exposure of worms to blue light increases the locomotion of *glo-1* mutant worms that are defective in gut granule biogenesis to a similar extent as wild-type worms (Figure S1). These results suggest that gut granules are not involved in the worms’ light avoidance behavior.

### Autofluorescent granules are widespread in nematodes

To explore whether the biology behind the green flash phenomenon is common in nematodes, we first examined whether gut granule-like autofluorescent granules are present across different nematode species.

Gut granules in *C. elegans* are considered to be rhabditin granules (Laufer et al. 1980). Birefringent granules in “*Rhabditis”* species were first described by Maupas (1900), with the term “rhabditin” used to describe the organic material (crystal) that was the likely source of the birefringence (Cobb 1914; Chitwood and Chitwood 1950). Such birefringent granules are common in “*Rhabditis”* species (including the current nematode genus such as *Caenorhabditis*, *Oscheius*, and *Pelodera*) (Cobb 1914; Chitwood and Chitwood 1950; Maupas 1900; Thomas and Quastler 1950; Bossinger and Schierenberg 1992), but have also been reported in an evolutionary distant nematode, *Spiroxys contortus* (Hedrick 1935), and autofluorescence has been reported to be widespread in nematode intestines in the form of globules (Forge and Macguidwin 1989).

We examine the intestinal autofluorescence of five other species of nematodes that are evolutionarily divergent but have established laboratory culture methods. These are *Oscheius tipulae* (Lam and Webster 1971); *Pristionchus pacificus* (Sommer et al. 1996); *Steinernema hermaphroditum* (Stock et al. 2004); *Panagrellus redivivus* (Linnaeus, 1767), and a species of the *Panagrolaimus* genus (*Panagrolaimus sp.* PS1159) (Sommer and Sternberg 1996). With *C. elegans* included, we examined three members each of Clade IV (*Steinernema*, *Panagrellus*, *Panagrolaimus*) and V (*Caenorhabditis*, *Oscheius*, *Pristionchus*) nematodes based on the five major clade classification of Blaxter et al. (1998).

In all species examined, we found autofluorescent granules to be present at least in the early and late larval stages (Figure 4), with observable differences in distribution and morphology. In *O. tipulae* and *P. redivivus*, some of the granules are strikingly birefringent and can easily be observed with Nomarski differential interference contrast microscopy (DIC) (Figure 5). At higher magnification, at least some of these birefringent granules have a Maltese cross appearance (Figure 5A, D), similar to what has been described by Cobb (1914). The birefringent granules are also autofluorescent (Figure 5B), but at least under Nomarski microscopy (DIC), not all autofluorescent granules are birefringent.

**Figure 4.**
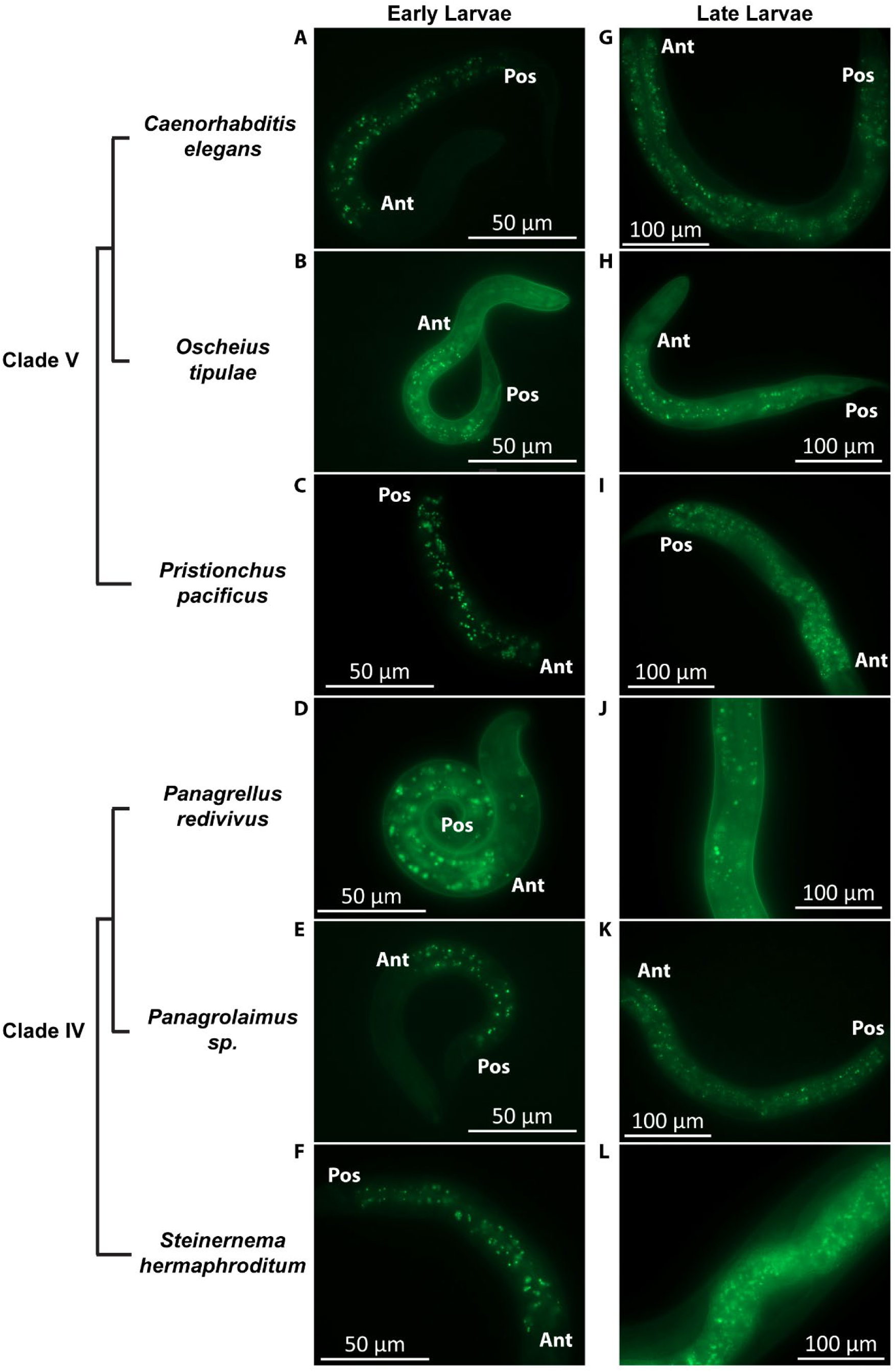
Autofluorescent granules are widespread in nematodes. Epifluorescence images (eGFP channel) of (A-F) Early Larvae. (G-L) Late Larvae. (A) L1 *Caenorhabditis elegans*. (B) L1 *Oscheius tipulae*. (C) J2 *Pristionchus pacificus*. (D) L1 *Panagrellus redivivus*. (E) L1 *Panagrolaimus sp.* PS1159. (F) J1 *Steinernema hermaphroditum*. (G). L4 *C. elegans*. (H) L4 *O. tipulae*. (I) J4 *P. pacificus*. (J) L4 *P. redivivus*. (K) L4 *Panagrolaimus sp.* PS1159. (L) J4 *S. hermaphroditum*. “Ant” indicate the anterior end of the intestine; “Pos” indicate the posterior end of the intestine. Scale bar: (A-F): 50µm; (G-L): 100µm.

**Figure 5.**
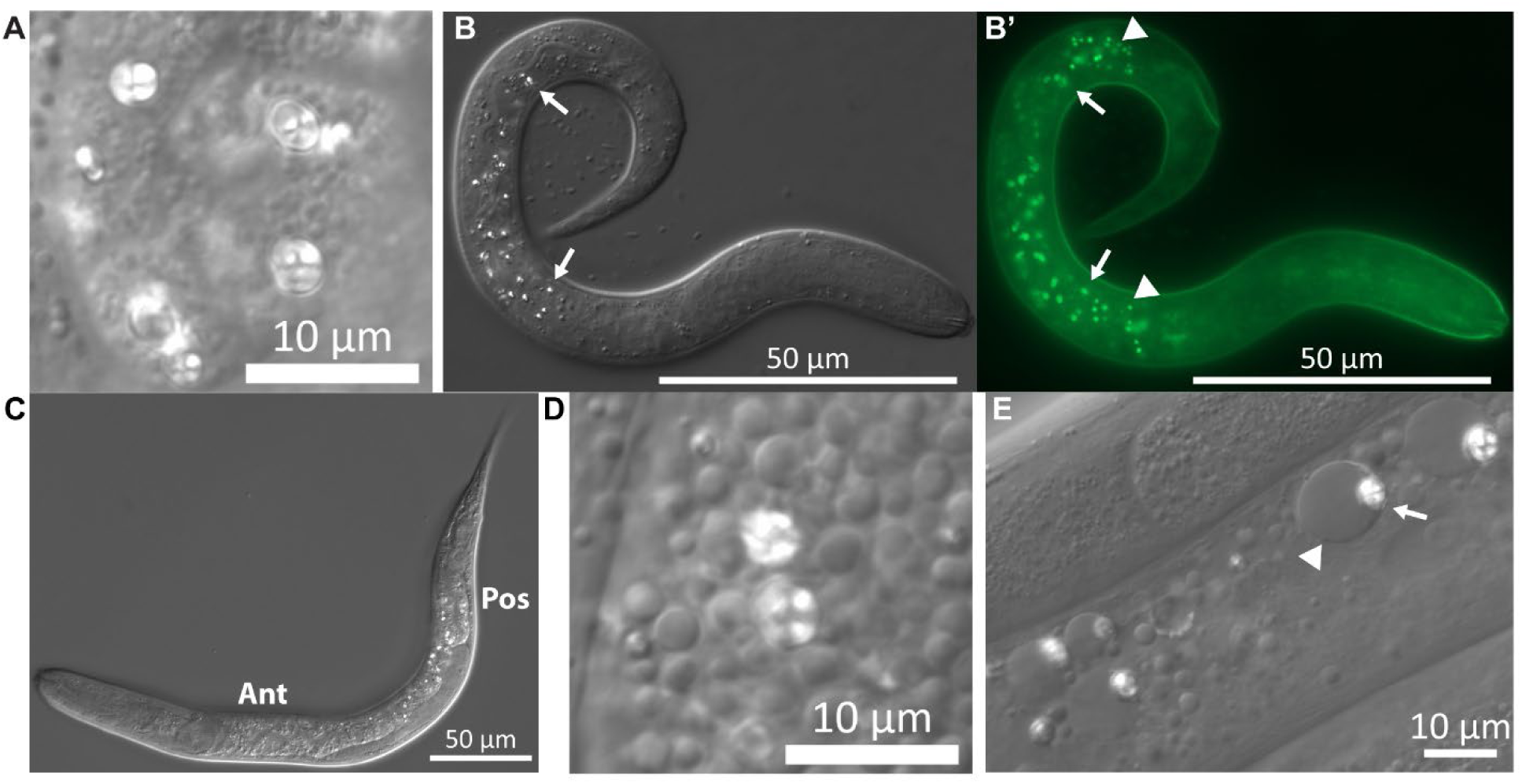
Birefringent granules in nematode intestine. Birefringent granules can be easily observed with Nomarski microscopy (DIC) in the intestine cells of *O. tipulae* (A-C) and *P. redivivus* (D-E). (A) Birefringent granules of *O. tipulae*. Some granules have a Maltese cross appearance. The image is from an adult hermaphrodite. (B-B’) A L1 *O. tipulae* larvae imaged with DIC (B) or eGFP channel (B’). The birefringent granules are also autofluorescent, but autofluorescent granules may not always be birefringent. Arrows point to two granules that are both birefringent and autofluorescent. Arrowheads point to two granules that are autofluorescent but not birefringent. Most birefringent granules are in the mid-gut, while most of the autofluorescent granules in both ends of the intestine are not birefringent. The head of the animal is toward the right. (C)A mid-larvae (∼L3) stage *O. tipulae* imaged with DIC. The distribution of the birefringent granules shows a strong bias towards the posterior half of the intestine. “Ant” indicates the anterior end of the intestine; “Pos” indicates the posterior end of the intestine. (D) Birefringent granules of *P. redivivus*. Some granules have a Maltese cross appearance. The image is from an L4 female. (E) Birefringent granules (arrows) in the intestine of adult *P. redivivus* are often associated with larger non-birefringent granules (arrowhead). The granules are closely associated, as can be seen in Video S5. Scale bar: (A, D, E): 10 µm; (B, C): 50 µm.

In *O. tipulae*, the pattern in which these birefringent granules are distributed in the intestine varies by stage (Figure 5B, C). In J1 larvae, they are predominantly mid-gut (Figure 5B), but by later stages, a clear spatial difference emerges (Figure 5C). Specifically, the autofluorescent granules at both ends of the intestine remain largely non-birefringent throughout development (Figure 5B, 6A, B, C, S2). The birefringent granules and non-birefringent autofluorescent granules might represent two distinct groups of LROs.

In adult *P. redivivus*, the birefringent granules are often clearly associated with a larger non-birefringent granule (Figure 5E, Video S5). The association bears a resemblance to the bilobed structure in the gut granules of *C. elegans* observed under excessive and deficient zinc conditions (Roh et al. 2012; Mendoza et al. 2024), with the larger non-birefringent granule resembling the expansion compartment observed.

### Gut granule green flash is common in nematodes

After establishing that autofluorescent granules similar to gut granules are common in nematodes, we explored whether this is also true for the “green flash” phenomenon. Of the six species (including *C. elegans*) tested, we were able to reliably observe and record the phenomenon in all but *Panagrolaimus sp.* PS1159 (Table 1, Figure 6, S2, Video S7-11). In all species with the green flash phenomenon, it could always be observed in at least some of the larvae/ juvenile stage four (L4/J4) animals (Late larvae) (Figure 6F-M, S2A-B, Video S8-10). This commonality persists despite the large difference in size, morphology, and lifestyle of these species, suggesting that there might be some common biological process that occurs at this stage of development. In addition to the L2 and dauer stage *C. elegans* (Figure 1A-E, Video S1), we observed the phenomenon in the J1 animal of *S. hermaphroditum* and the adult (hermaphrodite) animal of *O. tipulae* (Figure 6A-E, S2C-E, Video S7,11), so it is not exclusive to late larval stages (Table 1).

**Figure 6.**
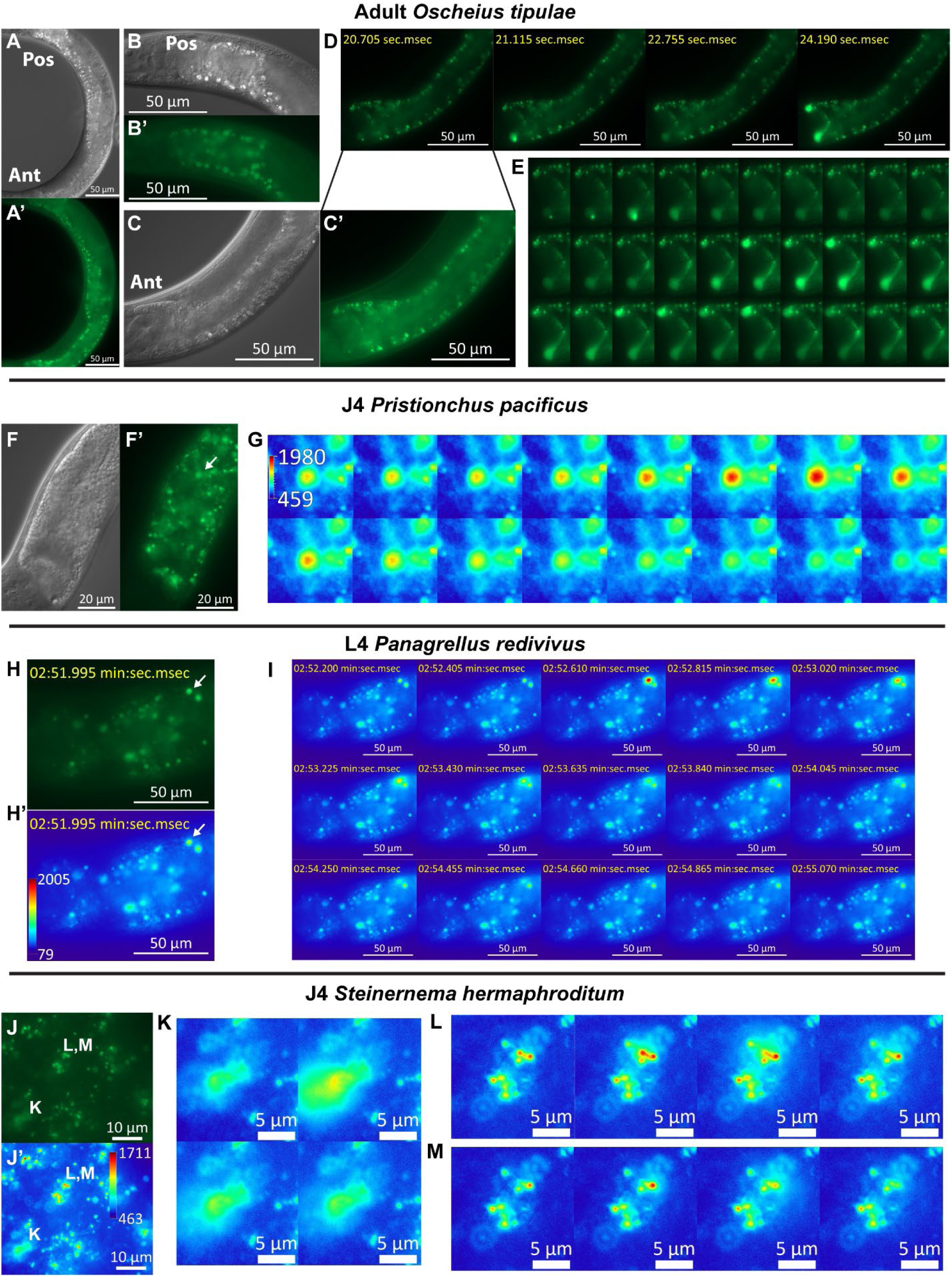
The green flash phenomenon of autofluorescent granules in intestinal cells are shared among nematodes. (A-E) Adult hermaphrodite *O. tipulae*. Scale bar: 50 µm (A) Nomarski microscopy (DIC) and (A’) epifluorescence (eGFP) image of the worm’s intestine. “Ant” indicates the anterior end of the intestine; “Pos” indicates the posterior end of the intestine. The distribution of the birefringent granules has a strong posterior bias. (B-B’) The enlarged DIC (B) and eGFP (B’) image of the posterior end of the intestine. No obvious green flash phenomenon was observed in this region (Video S6).(C-C’) The enlarged DIC (C) and eGFP (C’) image of the anterior end of the intestine. The autofluorescent granules at the anterior end are not birefringent under DIC and are often somewhat smaller in size. (D) Fluorescence images (eGFP channel) of the anterior end of the intestine at different time points during the analysis. Values represented are seconds and milliseconds and start (from 0) with the initiation of live imaging. The phenomenon is most commonly observed in the first intestinal cell ring. The complete series of images can be found in Video S7. (E) eGFP time series focusing on the first intestinal cell ring, covering largely the same period as shown in D (start at the same frame), the images were acquired at a rate of ∼5 frames per second. (F-G) J4 hermaphrodite *P. pacificus*. (F-F’) Nomarski microscopy (DIC) (F) and epifluorescence (eGFP) (F’) image of the anterior intestine. The arrow points to the gut granule detailed in the time series (K). Scale bar: 20µm. (K) eGFP (pseudo-colored-heat map) time series of the green flash of the granule indicated by the arrow in F’, the images were acquired at a rate of ∼5 frames per second. The number on the vertical bar indicates the range of the color assignment of the heat map. The complete series of images can be found in Video S8. (H-I) epifluorescence (eGFP) images of the anterior intestine of a J4 female *P. redivivus*. Scale bar: 20µm. (H-H’) pseudo-colored-green (H) and pseudo-colored-heat map (H’). The arrow points to a gut granule that flashes, as shown in the time series (I). Scale bar: 50µm. The number on the vertical bar (H’) indicates the range of the color assignment of the heat map. (I) Time series: The images were acquired at a rate of ∼5 frames per second. Scale bar: 50µm. (J-M) epifluorescence (eGFP) images of a section of the anterior intestine of a J4 hermaphrodite *S. hermaphroditum*. pseudo-colored-green (J) and pseudo-colored-heat map (J’). The labels “L, M” and “K” indicate the corresponding enlarged time series images in K and “L, M” (L and M are two time series of the same region). Scale bar: 10µm. The number on the vertical bar (J’) indicates the range of the color assignment of the heat map. Scale bar: 5 µm. The complete series of images with lower resolution can be found in Video S9.

**Table 1.**
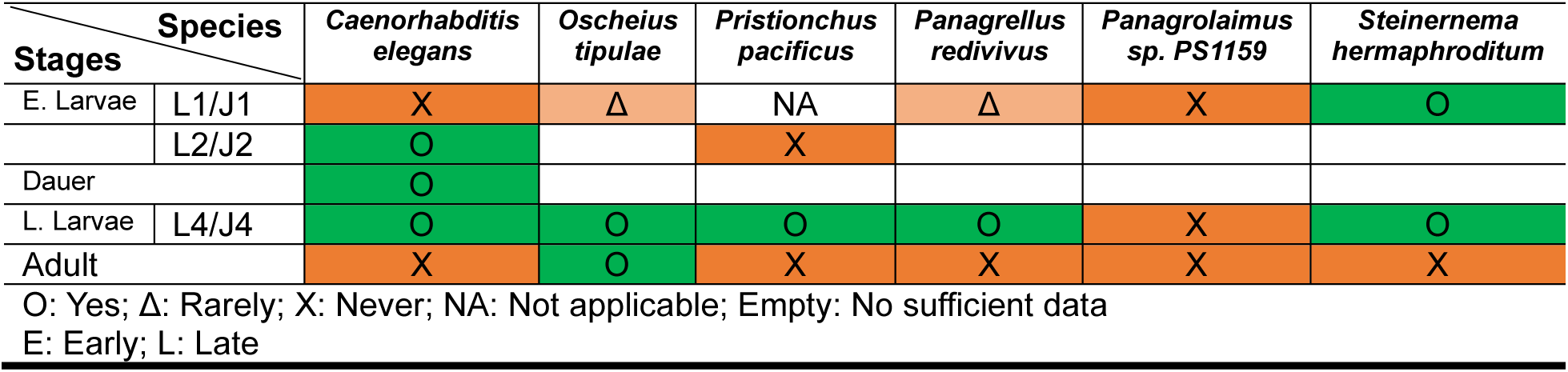
Developm ental stages in which gut granule green flash can be observed.

In both late larvae and adult animals, we also observed that there is a strong anterior bias for the green flash phenomenon. When observed, it occurred at the anterior end of the intestine, but rarely at the posterior end. For example, in both J4 hermaphrodite *S. hermaphroditum*, and adult hermaphrodite *O. tipulae,* green flash can be induced at the anterior end of the intestine in all animals (six of six each) with the phenomenon (Figure 6C-E, J-M, Video S7, S9), but none at the posterior end (Figure 6B, Video S6); in J4 hermaphrodite *O. tipulae*, green flash can be induced at the anterior end of the intestine in all ten animals (Figure S2 A-B, Video S10), but only weakly at the very end of the posterior end in 4 of 10 animals. At least in *O. tipulae*, the phenomenon also occurs predominantly at the ends of the intestine, with the strongest intensity observed at the first anterior intestinal cell ring (Figure 6D-E, S2A-B, Video S7, S10). It is worth noting that the distribution pattern of the flashable autofluorescent granules in *O. tipulae* (anterior-biased, concentrated at the ends of the intestine) is opposite to the autofluorescent granules that are also birefringent, as previously noted (posterior-biased, concentrated at the middle section of the intestine). In both *O. tipulae* and *P. redivivus*, we did not observe obvious green flash events in the birefringent granules.

Overall, we conclude that the population of autofluorescent granules in nematodes comprises two or more subpopulations with distinctive differences in biochemical properties that undergo significant changes during development, and that the green flash phenomenon is one of the assays that could be used for identifying these populations.

## Discussion

*C. elegans* has long been known to be autofluorescent (Babu 1974), with most of the autofluorescence coming from the gut granules (Clokey and Jacobson 1986). Particular interest has been paid to the increasing level of autofluorescence as the worms age or are stressed (Klass 1977; Davis et al. 1982; Clokey and Jacobson 1986; Forge and Macguidwin 1989; Pincus et al. 2016; Hajdu et al. 2023). However, the fluorophores responsible for gut granule autofluorescence remain largely undefined. One of the fluorophores emitting blue fluorescence has been identified as anthranilic acid (Babu 1974; Coburn et al. 2013), and changes in the blue fluorescence occur near the death of the animals (<u>d</u>eath fluorescence) have been described (Coburn et al. 2013; Pincus et al. 2016). Both green and red fluorescence also increase with age, and the green fluorescence intensifies near the death of the animals (Coburn et al. 2013; Pincus et al. 2016). It is important to note that these fluorescence changes are distinct from the gut granule green flash phenomenon that we are describing in this research, both spatially and temporally.

In *C. elegans*, gut granules have been shown to have both metabolic and signaling functions. These lysosome-related organelles (LROs) are a site for fat, cholesterol, and heme storage (Schroeder et al. 2007; Lee et al. 2015; Sun et al. 2022) as well as a storage and sequestering site for micronutrient metals (Roh et al. 2012; Chun et al. 2017). Gut granules are also the site of biosynthesis for secondary metabolites, including the signaling molecules ascarosides (Le et al. 2020). Recently, they have also been suggested to be involved in the responses to other forms of stress and in the resistance to pathogens (Hajdu et al. 2023). They are also structurally more complex than a simple sphere with a lipid bilayer. The studies of Roh et al. (2012) and Mendoza et al. (2024) show that gut granules are bilobed with two compartments, which they describe as an acidified compartment and an expendable compartment. The structure is also not static, with the volume of the expendable compartment changing at least in response to dietary zinc levels (Roh et al. 2012; Mendoza et al. 2024). The mechanism of such a structural rearrangement remains largely uncharacterized. Unexpectedly, in our survey of nematode intestine granules, we discovered a striking similarity in the granules of *P. redivivus* (Figure 5E), an evolutionary distant nematode, suggesting that such organelle structure is not unique to *C. elegans* and might be common in nematodes and maybe beyond.

In an attempt to develop the intestine of *C. elegans* as a model to study vesicle release, we discovered a novel phenomenon of dynamic autofluorescence in intestinal lysosome-related organelles. When observed with the green fluorescence channel, these autofluorescent granules of the nematode intestine underwent a rapid and dynamic change in intensity, a phenomenon we describe as “gut granule green flash.” Observable changes begin with a sharp increase in fluorescence intensity and end with rapid dissipation and diffusion. Further analysis suggests that at least two distinctive fluorophores or fluorophore transitions exist in the gut granules, with only one of them having the “flash” characteristic.

Mechanistically, the green flash phenomenon could be explained either by the breakdown of the granule and the release of one or more of its constituents (Model 1, Figure 7A), or the release of some of the granule constituents by granules that remain intact (Model 2, Figure 7B). In both models, changes in the green fluorescence intensity may be explained by the green fluorophore responding to differences between the lysosome-related organelles and the cytoplasm. The lumens of the lysosome-related organelles are more acidic than the cytoplasm (Clokey and Jacobson 1986; Kostich et al. 2000; Hermann et al. 2005); If the green fluorophores are pH-sensitive, with fluorescence intensity positively correlated with pH, the green fluorophore would increase in fluorescence upon exiting the granules. On the other hand, the changes in red fluorescence intensity may be explained by the red fluorophore being pH-neutral. The subsequent dissipation and diffusion can also be reasonably explained as the diffusion of both of these molecules. We think Model 1 may be more likely (Figure 7A), partly because there is some similarity between our observation and that of Luke et al. (2007), in which fluorescently labeled gut granule disruption in osmotic sensitive mutants under hypotonic shock was shown. Also, although the green flash phenomenon is distinct from the death (blue) fluorescence described (Coburn et al. 2013), as the green flash phenomenon is local (surrounding the LRO) and fast, and the death (blue) fluorescence is global and slow, two observations from different nematode species lead us to suspect that both could be caused by the breakdown of the gut granules. First, we found that at least in the dauer stage of *C. elegans,* continuous exposure to the shorter wavelength DAPI channel quickly turned the intestine/worm blue (Video S4). The change in the blue fluorescence bears some resemblance to the death (blue) fluorescence described. Second, although it is unclear for most of the nematode/ stages we analyzed, the gut granule green flash does seem to contribute to the build-up of overall cytostome green fluorescence in the anterior intestine of *O. tipulae* (Video S10). We do not exclude Model 2, however, as we noticed that it is not uncommon for residual autofluorescence to remain after the flash event for individual granules (Figure 1E), and in the *ex-vivo* confocal recorded experiments, we have also observed multiple flash events in some of the granules, both of which argued against the disruption of the granules and support substance release by intact granules as described in Model 2 (Figure 7B). It is also possible that the event described in both models occurs, but in different granules.

**Figure 7.**
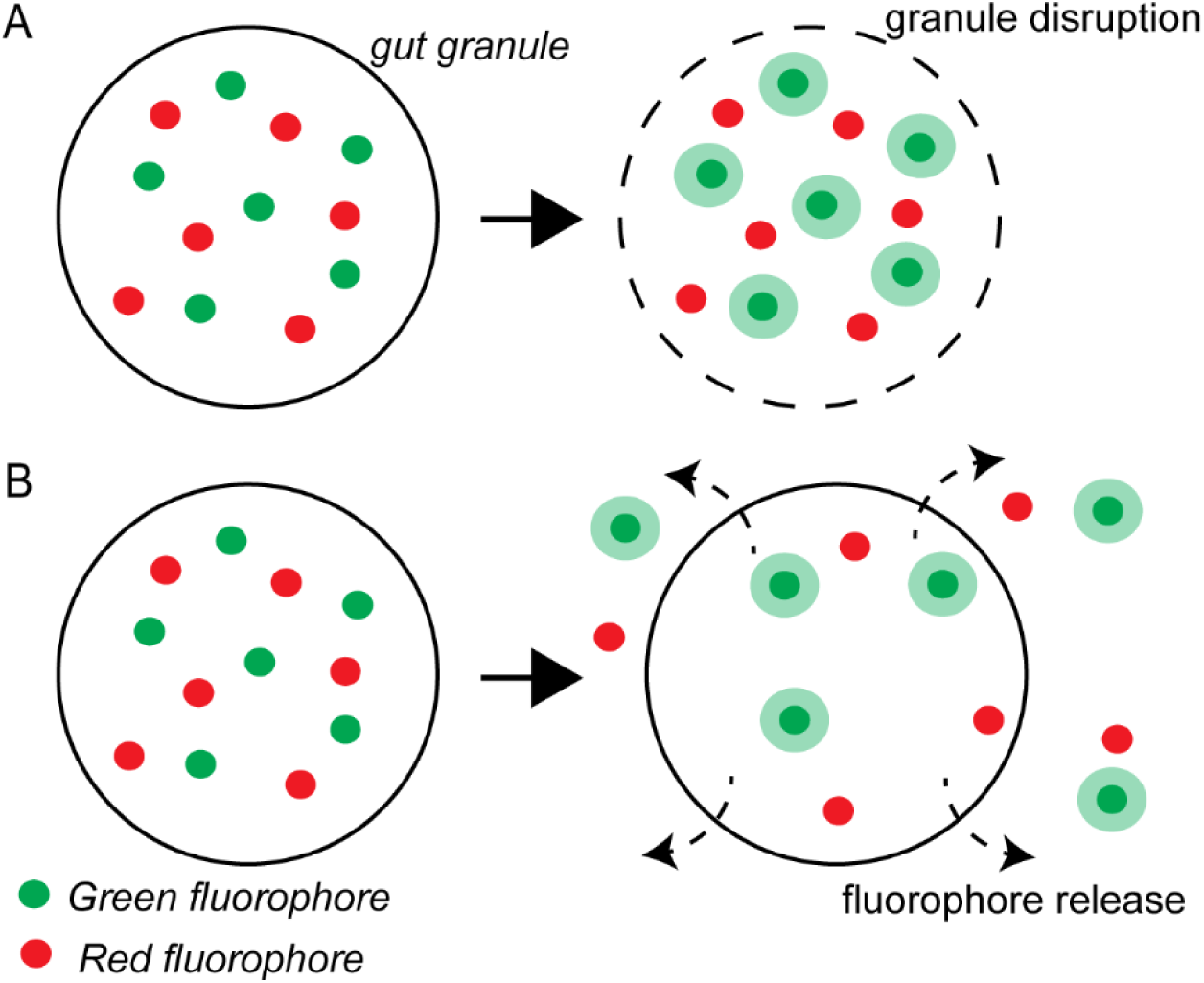
Models for the gut granule green flash phenomenon. In both models, the green fluorophores are pH-sensitive, with fluorescence intensity positively correlated with pH. On the other hand, the red fluorophores are likely pH-neutral. A) In the first model, the fluorescence dynamic is explained by the breakdown of the granule and the release of its constituents. The fluorescence of the green fluorophores increases after breakdown due to the higher pH environment in which the fluorophores now reside. B) In the second model, the fluorescence dynamic is explained by the release of the fluorophores by intact granules. The released green fluorophores increased in fluorescence in the more basic cytosol.

Regardless of the potential mechanisms, we believe that stimulation via blue light is likely the cause, as the phenomenon was usually not observed immediately upon illumination, and that increasing the intensity of the blue light significantly shortens the amount of time it takes for the flash to occur. However, we cannot rule out that gut granule green flash may also be naturally occurring. In the *ex-vivo* setting, the phenomenon was highly associated with the immediate presence of food (Figure 2), which may suggest that the phenomenon can be a part of or a consequence of a metabolic or signaling process in food intake. One possibility is that this phenomenon is associated with the release of vesicle content from gut granules, and the fluorophores or the content released play metabolic or signaling roles in food uptake. Alternatively, the exposure of the basal face of intestinal cells to bacteria and their metabolites could be a significant stressor to the cells, and it is plausible that the physiological condition in these cells caused the granules to be in a more “flashable” state.

We observed that not all gut granules can be induced to flash in our settings, and their inducibility varies significantly during development. We suspect that this is due to the existence of multiple sub-groups of gut granules and that their biology changes significantly in the developmental process. To put the observed phenomenon in a wider context and to rule out that the stage-specific inducibility may be associated with worm size or something specific to *C. elegans*, we explored the green flash phenomenon in other nematode species. We found that the understanding of similar granules in other nematodes is somewhat limited. The first documented description of such granules (described as birefringent, blackish, and opaque with transmitted light) may be that of Maupas (1900), in the same research that also first describes *C. elegans* (*Rhabditis elegans*) as a species. Maupas (1900) suggests the lack of those granules in *C. elegans*. With better microscopy, it is now known that at least some (Laufer et al. 1980), if not all (Hermann et al. 2005), gut granules in *C. elegans* are clearly birefringent. Cobb (1914) used the term “rhabditin” to describe the organic material (crystal) that was the likely source of the birefringence he identified in the intestinal cells of *Mesorhabditis monhystera* (*Rhabditis monhystera*), and such “rhabditin granules” have since been found to be common among “*Rhabditis”* species (including the current nematode genus such as *Caenorhabditis* and *Oscheius*) (Cobb 1914; Chitwood and Chitwood 1950; Maupas 1900; Thomas and Quastler 1950). Similar granules have been described outside of the “*Rhabditis”* species; Maupas (1900) described two species that now belong to the genus *Pristionchus* with birefringent intestinal granules/corpuscles, and Hedrick (1935) used “rhabditin” to describe a granular substance that the author observed in an evolutionarily distant nematode, *Spiroxys contortus*. The autofluorescent nature of the granules was described much later (Laufer et al. 1980; Hermann et al. 2005), and mostly only in *C. elegans*, but Forge and Macguidwin (1989) found autofluorescence “globules” in all 15 diverse species of nematodes they analyzed, and Bossinger and Schierenberg (1992) described the autofluorescence pattern of gut primordium in *Oscheius dolichura* (*Rhabditis dolichura*, also described by (Maupas 1900) to be birefringent) embryos in comparison to that of *C. elegans*. It is likely that, in most of these species, the birefringent or autofluorescent granules/corpuscles/globules correspond to the gut granules of *C. elegans*.

We found autofluorescent granules to be common in all nematode species we analyzed. Other than *C. elegans*, we examined *Oscheius tipulae* (another *Rhabditis*), *Pristionchus pacificus* (an evolutionarily distant Clade V (Suborder Rhabditina) species), and three Clade IV (Suborder Tylenchina) species, namely *Steinernema hermaphroditum*, *Panagrellus redivivus*, and *Panagrolaimus sp.* PS1159. In our recent estimation, the divergent time of the Rhabditina and Tylenchina species was estimated to be over 200 million years (Schwarz et al. 2025). We showed that many granules of *O. tipulae* and *P. redivivus* were highly birefringent under Nomarski DIC optics, which rely on circularly polarized light (Figure 5). In both *O. tipulae* and *P. redivivus,* the appearance of the birefringent granules closely matched the sketches of Cobb (1914). In *O. tipulae*, the distribution of the birefringent granules depends on the stage of the animal and, at least in later stages, has a clear posterior bias. Maupas (1900) reported a similar posterior bias in several of the nematode species he analyzed, including a species now belonging to the *Oscheius* genus (*O. dolichura*, originally *Rhabditis dolichura*). The presence of autofluorescent granules across species and the birefringent granules in *P. redivivus* suggests that “rhabditin granules” are likely to be widespread among nematodes beyond “*Rhabditis*” species.

Analyzing the selected worms, we found that in addition to *C. elegans*, the green flash phenomenon can be observed in at least some stages of *O. tipulae*, *P. pacificus*, *P. redivivus*, and *S. hermaphroditum* (Table 1). In particular, in all five species, the phenomenon can be observed during the L4/J4 stage despite the significant difference in the size and shape of these animals, suggesting that the susceptibility of these granules is likely due to their stage-specific biochemical properties and not to factors such as the thickness of the sample (worm). As in *C. elegans*, we found that the susceptibility of the granules differs within each animal, and at least in some species, we found this to be associated with their overall location within the intestine and with other characteristics of the granules, such as the lack of easily observable birefringence. Together, these findings suggest different sub-populations of gut granules in the intestine of the nematodes. This is also the conclusion from another recent independent study of autofluorescence of these granules in *C. elegans* (Hulsey-Vincent et al. 2024).

Further understanding of gut granule green flash may require the characterization of the green and red fluorophores. It is unclear how to best identify these fluorophores, although we would predict that the green fluorophores emit stronger fluorescence under high pH conditions, while the red fluorophores are likely pH neutral. Whether or not the phenomenon is associated with the presence of the corresponding fluorophores is also uncertain. Regardless, one plausible approach to the biochemical causes of the phenomenon would be to compare the molecular content of wild-type and *glo-1(lf)* mutants that lack gut granules, potentially with mass spectrometry, or with fluorescence spectroscopy. Since our results identify the stages in which the green flash can be induced across different species, stage and species-specific samples that contain or lack gut granules could be used to narrow down the candidates. In the non-*C. elegans* species that we analyzed, CRISPR-Cas9 genome modification methods are technically available for *O. tipulae* (Dockendorff et al. 2022), *P. pacificus* (Witte et al. 2015)*, S. hermaphroditum* (Schwartz et al. 2024; Cao 2023), and *Panagrolaimus sp.* PS1159 (Hellekes et al. 2023), so knocking out the *glo-1* orthologs may be an effective way to eliminate gut granules in those species. Similarly, one could use proteomics, as we recently reported: we identified gut granule proteins by comparing the proteome of the *C. elegans* wild-type intestine to that of the *glo-1* mutants (Tan et al. 2024).

Overall, we described a visually spectacular fluorescence dynamic phenomenon that involves a type of lysosome-related organelle known as gut granules, which we showed to be common in nematode intestines. We also found that gut granules are likely a collection of distinct subgroups that change during development, and the dynamic fluorescence phenomenon we describe might provide a handle on their eventual characterization.

## Materials and Methods

### Nematode strain maintenance and general methods

The culture and maintenance of *C. elegans, O. tipulae*, *P. pacificus*, *P. redivivus*, and *Panagrolaimus sp.* PS1159 was done similarly to the standard *C. elegans* procedures as described in Brenner (1974). Briefly, worms were cultured on 6 cm Petri dishes containing Nematode Growth Medium (NGM) agar with a bacterial lawn of *Escherichia coli* strain OP50. *C. elegans, O. tipulae*, *P. pacificus*, and *P. redivivus* were maintained at 20°C, and *Panagrolaimus sp.* PS1159 at 25°C. The culture and maintenance of *S. hermaphroditum* were done similarly to the procedures described in Cao et al. (2022) and Rodak et al. (2024). Briefly, worms were either cultured on 10cm Petri dishes containing NGM agar with a bacterial lawn of *Xenorhabdus griffiniae* strain HGB2511 or 10cm Petri dishes containing Enriched Peptone Medium (EPM) with a bacterial lawn of *Comamonas aquatica* DA1877 (Avery and Shtonda 2003) at 25°C. For *C. elegans*, the Bristol N2 strain (Brenner 1974) was used as the wild-type reference strain and is the strain from which the *glo-1* mutant strain was derived. The five times out-crossed *glo-1(zu391*lf) X (Hermann et al. 2005) mutant strain OJ1347 (Wang et al. 2013) was a gift from Dr. Derek Sieburth of the University of Southern California. For *O. tipulae*, *P. pacificus*, *S. hermaphroditum*, *P. redivivus*, and *Panagrolaimus sp.*, the following wild-type reference strains were used, respectively: CEW1 (Winter 1992), PS312 (Sommer et al. 1996), PS9179 (Cao et al. 2022), MT8872 (Sternberg and Horvitz 1981), and PS1159 (Sommer and Sternberg 1996).

### Preparation of *C. elegans* dauer larvae

To induce dauers, 10-20 young adults were placed on 35 mm diameter Petri dishes containing 2 mL of NGM agar (without peptone) supplemented with a quantity of crude pheromone extract (Schroeder and Flatt 2014) that normally induced 95-100% of dauers in wild-type animals – typically 10-30 µL per 2 mL of agar. Plates were seeded with 10 µL of 8% w/v *E. coli* OP50 that had been heat-killed at 95°C for 5 minutes. Adults were picked onto the plate and allowed to lay eggs at room temperature (RT; 22-23°C) for 5-9 hours before being removed, during which time they typically laid 200-300 eggs. The plates were then further seeded with an additional 20-50 µL of heat-killed OP50, then wrapped with Parafilm and incubated at 25.5°C for 60-72 hours. Dauers were then picked from this plate for analysis.

### *in-vivo* imaging and analysis of gut granule autofluorescence dynamics

*in-vivo* imaging and analysis were performed with a Zeiss Imager Z2 microscope equipped with an Apotome 2 and Axiocam 506 mono using Zen 2 Blue software. Worms were partially immobilized with 1 mM levamisole in S-basal medium (Stiernagle 2006) and mounted on 2 or 5% agarose pads on microscope slides and observed with a 100X objective.

The excitation wavelength (filter) was 450-490nm, and the emission filter was 500-550 nm. If no green flash phenomenon was observed within 180-300 s of continuous light exposure, the worm or intestinal region was characterized as not having the phenomenon. The observation was either done through live observation through the eyepiece of the microscope or by reviewing the acquired image series. For the DAPI channel experiments, the excitation wavelength (filter) was 370-410 nm, and the emission filter was 430-470 nm. Image series (or video) were acquired at a rate of about 5 frames per second, except for the experiments with L4 stage *C. elegans* larvae, which were acquired at a rate of about 6.5 frames per second.

Hermaphrodite or female adult and late larvae (L4/J4) animals were selected based on vulva and gonad morphology. In *C. elegans*, adult animals were young adults developed from L4 animals selected on the previous day. First larval (juvenile) stage (L1/J1) animals of *S. hermaphroditum, O. tipulae*, *P. redivivus*, and *Panagrolaimus* sp. PS1159, and second juvenile stage (J2) animals of *P. pacificus* (*P. pacificus* hatches as J2(Fürst von Lieven 2005)) were selected based on size, general morphology, and gonad morphology (4-cell gonad). In *C. elegans,* first larval stage (L1) animals were obtained by hatching eggs (embryos) obtained by dissolving gravid hermaphrodites (using a mixture of bleach and sodium hydroxide) overnight in M9 buffer, similar to what was described in (Stiernagle 2006). Second larval stage (L2) *C. elegans* were obtained by allowing the synchronized L1 to develop for 24 hours at 20°C on NGM plates seeded with OP50.

### Sample preparation for *ex-vivo* imaging

L4 stage hermaphrodite worms were transferred to a drop of Iwasaki–Teramoto (I–T) solution (Teramoto and Iwasaki 2006) [136mM NaCl, 9mM KCl, 1mM CaCl_2_, 3mM MgCl_2_, 77mM D-glucose, and 5mM HEPES (pH 7.4)] and immobilized with 1mM levamisole on a microscopic slide. To expose the intestine for imaging, worms were incised with a pair of 30G 5/8” needles (PrecisionGlide, BD, Franklin Lakes, NJ) near the anus. For experiments with food added on the slide, a glob of OP50 collected by scraping the lawn off an NGM plate using a cell scraper was transferred into the droplet using a platinum-wire worm pick. A circle was drawn using a PAP pen (RPI) around the solution droplet to avoid spilling, and a piece of cover glass was subsequently mounted on top.

### Confocal Microscopy

Confocal time series were acquired using a Fluoview FV3000 Confocal laser scanning biological microscope with a 60×, 1.30 N.A. silicone oil objective (Olympus). Frame rate was calibrated to 1Hz by adjusting the size of the imaging window and the line averaging multiplier. Five minutes of time lapse images were acquired for each sample. All image processing and analyses were done using ImageJ (National Institutes of Health). The transmitted light channel was illuminated simultaneously with the imaging laser (473nm) for positional imaging. Panel A1 in Figures 4, as well as Panels B2, D2, F2 and G in Figure 3, were pseudo-colored with mpl-inferno LUT in FIJI (Schindelin et al. 2012).

### Confocal Imaging data processing

ROIs of granules were manually selected in FIJI for all data. For Figure 2(G-H), ROIs of peri-granule “cloud” were defined based on the identification of the fluorescent pattern of green flash events, as the surrounding rings that do not overlap with the granule. The zero points of the time series data were set at the phenomenon onsets of each granule and pooled into metadata.

ROI selection: ROIs were defined based on visual identification of granules in the intestine region, and the fluorescence normalization baseline was set accordingly as the average of the period prior to the changes in fluorescence. Standard deviation envelopes were visualized in the time series. For Figure 3(H), the number of gut granule green flash events was normalized as per minute *10^4^ µm^2^ of the intestinal surface area from single optical sections, which are manually delineated (Figure 3(G)). Nonparametric Mann-Whitney U test was used for comparison between WT (wild-type) food vs. no-food, and WT food vs. *glo-1(*lf).

### Light Stimulus and Locomotion Analysis

Locomotion analysis and animal size measurement were done similarly to what was described in Tan et al. (2022) and Wong et al. (2019) with adaptations. Briefly, at least an hour before the experiment, young adult worms (1 day post-L4) were transferred to NGM plates freshly seeded with 50µl of OP50. Five worms were transferred per plate. The recording and tracking of the worms were performed using WormLab (MBF Bioscience, Williston, VT) equipment and software. The camera was a Nikon AF Micro 60/2.8D with zoom magnification. Each plate was recorded for 2 minutes without stimulus followed by 5 minutes of blue light stimulus. The raw data from the WormLab was then manually processed by matching the tracking data with the recorded video. The average speed shown in Figure 5 was based on the average of the absolute value of the locomotion data (recorded at the frequency of 7.5Hz) collected at the time span of 2 seconds (15 frames).

### Data and statistical analysis

Data were plotted as mean ± s.e.m. in Figure 3 and as mean ± s.d. in Figure S1. In Figure 2I, as the raw data failed to pass all normality tests (Kolmogorov-Smirnov test, Shapiro-Wilk test, D’Agostino & Pearson test), the data were plotted as median ± 95% CI instead. Mann-Whitney U test was used in Figure 2 and Student’s *t*-test was used in Figure S1.

## Acknowledgments

We thank members of the Sternberg and Anderson laboratories for experimental support and discussions, particularly David J. Anderson, Han Wang, Xuan Wan, and Stephanie Nava, for their involvement in the earlier stages of this research. We would especially like to thank David J. Anderson and Hillel Schwartz for their valuable suggestions and critical reading of the manuscript. We would also like to thank Marie-Anne Felix for the English translation of the seminal work of Émile Maupas (Maupas 1900), which can be accessed on the Wormbase website (https://wormbase.org/, under Community/ Resources/ Key Papers). The five times out-crossed *glo-1(zu391 lf)* X (Hermann et al., 2005) mutant strain OJ1347 (Wang et al., 2013) was a gift from Derek Sieburth of the University of Southern California. This work was also facilitated by WormBase, a knowledgebase for nematode research (Sternberg et al. 2024), and Nemys: World Database of Nematodes (Nemys eds. 2025). This research was funded by NIH award UF1NS111697 and NSF Enabling Discovery through GEnomics (EDGE) grant 2128267 to PWS, a Bren Professor of Biology.

## Data Availability Statement

Strains are available upon request. Supplemental files available at FigShare.

---

**Video S1** Dynamic fluorescence intensities change in intestinal granules of a dauer stage *C. elegans* larva, pseudo-colored-heat map. Snapshots of this video are shown in Figure 1 (pseudo-colored-green). Gut granule green flash events can be seen starting at about 0:00:20 in this video. Values represented are seconds and milliseconds and start (from 0) with the initiation of live imaging. The number on the vertical bar indicates the range of the color assignment of the heat map; Scale bar: 10 µm.

**Video S2** Dynamic fluorescence intensities change in intestinal granules of an L4 stage *C. elegans*. Snapshots of this video are shown in Figure 1. Gut granule green flash events can be seen starting at about 0:01:20 in this video. Scale bar: 10 µm.

**Video S3** Dynamic fluorescence intensities change in intestinal granules of an L4 stage *C. elegans*. The video was taken with a scanning confocal microscope in the *ex-vivo* setting.

**Video S4** Continuous exposure to DAPI makes the whole worm/ intestine autofluorescent. Values represented are seconds and milliseconds, and start (from 0) with the initiation of live imaging.

**Video S5** Birefringent granules in the intestine of adult *P. redivivus* are tightly associated with larger non-birefringent granules. The granules always stayed together. Values represented are seconds and milliseconds, and start (from 0) with the initiation of live imaging. Scale bar: 50µm.

**Video S6** Gut granule green flash was not often observed in the posterior end of the nematode intestines. The video is of the posterior end of an adult *O. tipulae* intestine, the same animal and region shown in Figure 6B. Values represented are seconds and milliseconds, and start (from 0) with the initiation of live imaging.

**Video S7** Gut granule green flash in the anterior intestine of an adult hermaphrodite *O. tipulae*. The phenomenon is most commonly observed in the first intestinal cell ring. Snapshots of this video are shown in Figure 6D-E. Values represented are seconds and milliseconds, and start (from 0) with the initiation of live imaging. Scale bar: 50µm.

**Video S8** Gut granule green flash in the anterior intestine of a J4 hermaphrodite *P. pacificus*. Snapshots of a section of this video are shown in Figure 6G. Values represented are seconds and milliseconds, and start (from 0) with the initiation of live imaging. Scale bar: 20µm.

**Video S9** Gut granule green flash in the anterior intestine of a J4 hermaphrodite *S. hermaphroditum*. Snapshots of a section of this video are shown in Figure 6J-M. Scale bar: 10µm.

**Video S10** Gut granule green flash in the anterior intestine of an L4 hermaphrodite *O. tipulae*. The phenomenon is most commonly observed in the first intestinal cell ring. The green flash seems to contribute to the build-up of overall cell autofluorescence. Snapshots of this video are shown in Figure S2A-B. Values represented are seconds and milliseconds, and start (from 0) with the initiation of live imaging. Scale bar: 10µm.

**Video S11** Dynamic fluorescence intensities change in intestinal granules of an J1 stage *S. hermaphroditum*. Snapshots of this video are shown in Figure S2C-E. Gut granule green flash events can be seen starting at about 0:00:30 in this video.

**Figure S1.**
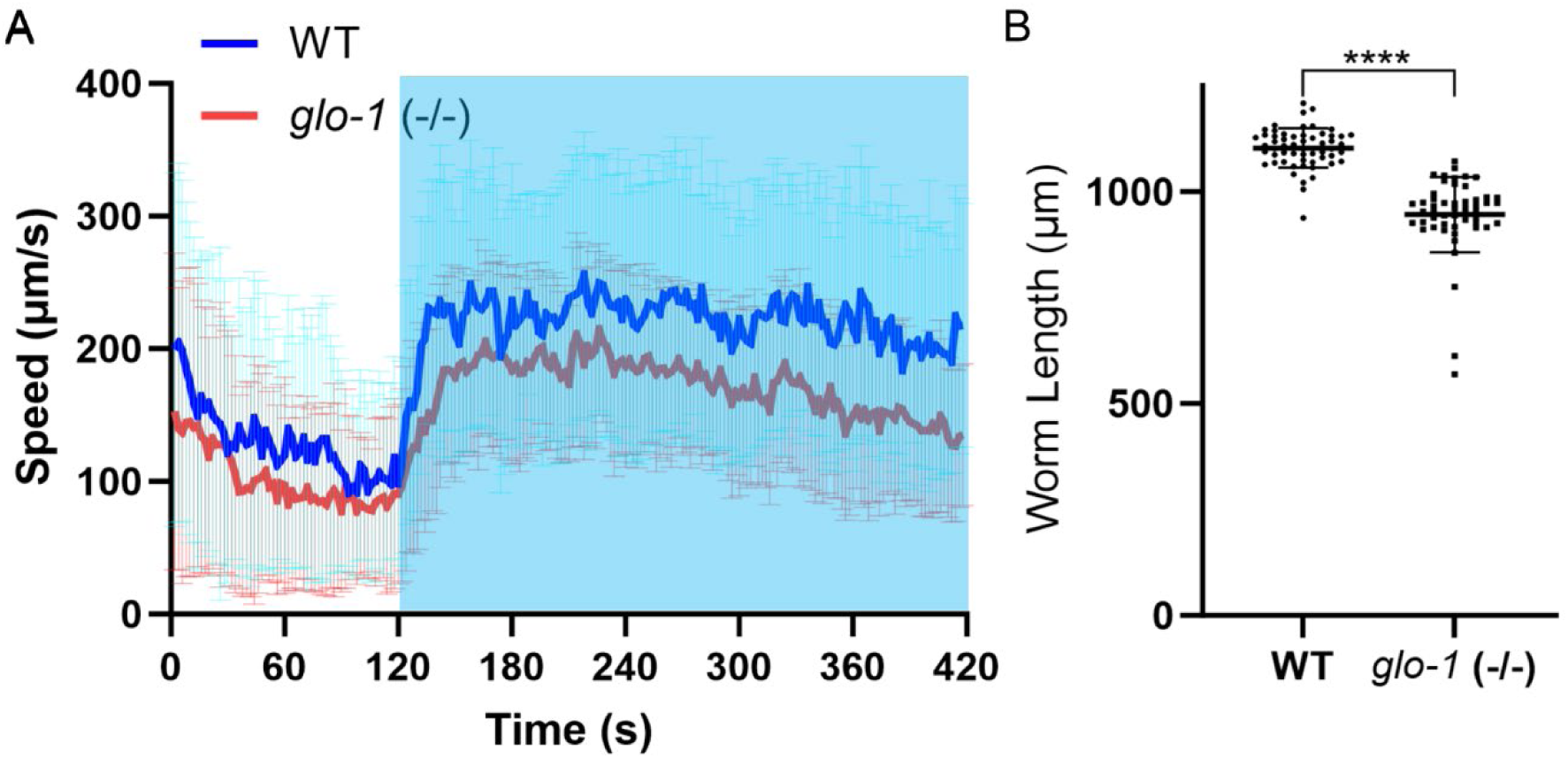
Gut granules do not play a significant role in the worms’ avoidance of blue light. (A) Both WT and *glo-1(zu391 lf)* worms respond to blue light applied between 120-420s (blue shade) by increased locomotion. Values are mean ± s.d (WT: n=57; *glo-1(zu391)*: n=52). *glo-1(zu391*lf) worms move slower with or without stimulus, which may partially be the result of having a smaller body size (B) *glo-1(zu391)* worms are slightly smaller compared to the wild type. Values are mean and error bars indicate Standard Deviation (WT: n=57; *glo-1(zu391)*: n=52) (****: p<0.0001, Student’s *t*-test).

**Figure S2.**
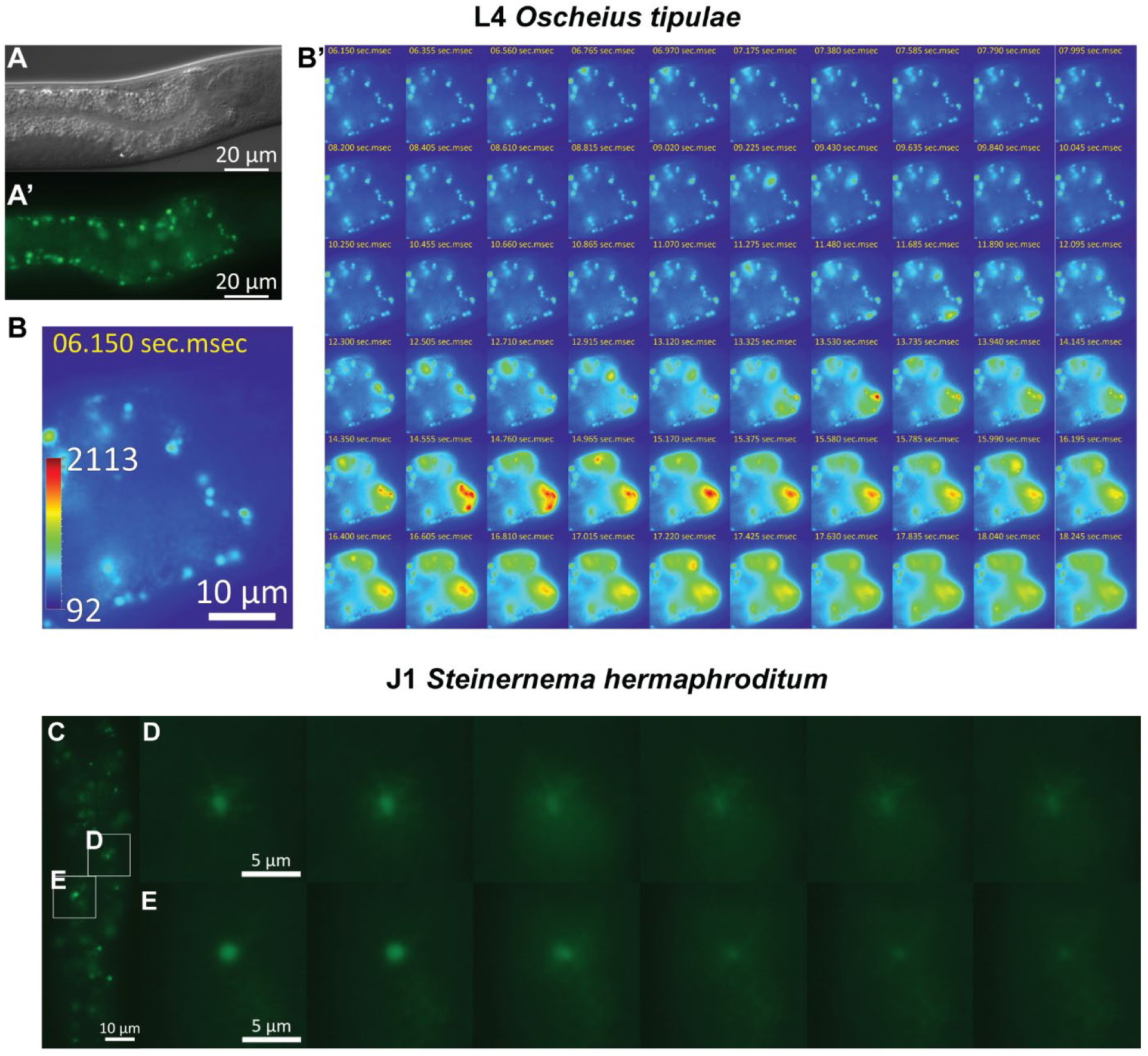
Gut granule green flash is common among nematodes. (A-B). The anteroior intestine of a L4 hermaphrodite *O. tipulae*. (A-A’) Nomarski microscopy (DIC) (A) and (A’) epifluorescence (eGFP) green (A’) image. Scale bar: 20 µm. (B-B’) The epifluorescence (eGFP) pseudo-colored-heat map image of the most anterior end of the intestine. The number on the vertical bar (B) indicates the range of the color assignment of the heat map. (B’) Time series showing the green flash phenomenon in the first cell ring, the images were acquired at a rate of ∼5 frames per second. The green flash seems to contribute to the build-up of overall cell autofluorescence. The complete series of images can be found in Video S10. Values represented are seconds and milliseconds and start (from 0) with the initiation of live imaging. Scale bar: 10 µm. (C-E) J1 *S. hermaphroditum.* (C) Epifluorescence images (eGFP) of the intestine of a J1 stage larva of wild-type *S. hermaphroditum*. Like those of *C. elegans*, the intestinal cells contain autofluorescent granules. Scale bar: 10 µm. (D, E) Enlargement of sections of the intestine shown in C, focusing on individual granules at the time of the dynamic changes over periods of ∼1 second. Scale bar: 5 µm. The complete series of images can be found in Video S5.

## References

1. Avery, L., and B.B. Shtonda, 2003 Food transport in the C. elegans pharynx. J Exp Biol 206 (Pt 14):2441–2457.

2. Babu, P., 1974 Biochemical Genetics of Coenorhabditis-Elegans. Molecular and General Genetics 135 (1):39–44.

3. Bhatla, N., and H.R. Horvitz, 2015 Light and hydrogen peroxide inhibit C. elegans Feeding through gustatory receptor orthologs and pharyngeal neurons. Neuron 85 (4):804–818.

4. Blaxter, M.L., P. De Ley, J.R. Garey, L.X. Liu, P. Scheldeman et al., 1998 A molecular evolutionary framework for the phylum Nematoda. Nature 392 (6671):71–75.

5. Bossinger, O., and E. Schierenberg, 1992 Transfer and tissue-specific accumulation of cytoplasmic components in embryos of Caenorhabditis elegans and Rhabditis dolichura: in vivo analysis with a low-cost signal enhancement device. Development 114 (2):317–330.

6. Brenner, S., 1974 The genetics of Caenorhabditis elegans. Genetics 77 (1):71–94.

7. Cao, M., 2023 CRISPR-Cas9 genome editing in Steinernema entomopathogenic nematodes. bioRxiv:2023.2011.2024.568619.

8. Cao, M., H.T. Schwartz, C.H. Tan, and P.W. Sternberg, 2022 The entomopathogenic nematode Steinernema hermaphroditum is a self-fertilizing hermaphrodite and a genetically tractable system for the study of parasitic and mutualistic symbiosis. Genetics 220 (1).

9. Chitwood, B., and M. Chitwood, 1950 Introduction to nematology (reprinted 1974). Baltimore: University Park Press.

10. Chun, H., A.K. Sharma, J. Lee, J. Chan, S. Jia et al., 2017 The Intestinal Copper Exporter CUA-1 Is Required for Systemic Copper Homeostasis in Caenorhabditis elegans. Journal of Biological Chemistry 292 (1):1–14.

11. Clokey, G.V., and L.A. Jacobson, 1986 The autofluorescent “lipofuscin granules” in the intestinal cells of Caenorhabditis elegans are secondary lysosomes. Mech Ageing Dev 35 (1):79–94.

12. Cobb, N.A., 1914 Rhabditin: contribution to a science of nematology. The Journal of Parasitology 1 (1):40–41.

13. Coburn, C., E. Allman, P. Mahanti, A. Benedetto, F. Cabreiro et al., 2013 Anthranilate fluorescence marks a calcium-propagated necrotic wave that promotes organismal death in C. elegans. PLoS Biol 11 (7):e1001613.

14. Dal Santo, P., M.A. Logan, A.D. Chisholm, and E.M. Jorgensen, 1999 The inositol trisphosphate receptor regulates a 50-second behavioral rhythm in C. elegans. Cell 98 (6):757–767.

15. Davis, B.O., Jr., G.L. Anderson, and D.B. Dusenbery, 1982 Total luminescence spectroscopy of fluorescence changes during aging in Caenorhabditis elegans. Biochemistry 21 (17):4089–4095.

16. Delevoye, C., M.S. Marks, and G. Raposo, 2019 Lysosome-related organelles as functional adaptations of the endolysosomal system. Curr Opin Cell Biol 59:147–158.

17. Dockendorff, T.C., B. Estrem, J. Reed, J.R. Simmons, S.B. Zadegan et al., 2022 The nematode Oscheius tipulae as a genetic model for programmed DNA elimination. Curr Biol 32 (23):5083–5098 e5086.

18. Edwards, S.L., N.K. Charlie, M.C. Milfort, B.S. Brown, C.N. Gravlin et al., 2008 A novel molecular solution for ultraviolet light detection in Caenorhabditis elegans. PLoS Biol 6 (8):e198.

19. Forge, T.A., and A.E. Macguidwin, 1989 Nematode autofluorescence and its use as an indicator of viability. J Nematol 21 (3):399–403.

20. Fürst von Lieven, A., 2005 The embryonic moult in diplogastrids (Nematoda) – homology of developmental stages and heterochrony as a prerequisite for morphological diversity. Zoologischer Anzeiger-A Journal of Comparative Zoology 244 (1):79–91.

21. Hajdu, G., M. Somogyvari, P. Csermely, and C. Soti, 2023 Lysosome-related organelles promote stress and immune responses in C. elegans. Commun Biol 6 (1):936.

22. Hedrick, L.R., 1935 Taxonomy of the nematode genus Spiroxys (family Spiruridae) … The life history and morphology of Spiroxys contortus (Rudolphi); Nematoda: Spiruruidae. Ann Arbor? Mich.

23. Hellekes, V., D. Claus, J. Seiler, F. Illner, P.H. Schiffer et al., 2023 CRISPR/Cas9 mediated gene editing in non-model nematode Panagrolaimus sp. PS1159. Front Genome Ed 5:1078359.

24. Hermann, G.J., L.K. Schroeder, C.A. Hieb, A.M. Kershner, B.M. Rabbitts et al., 2005 Genetic analysis of lysosomal trafficking in Caenorhabditis elegans. Mol Biol Cell 16 (7):3273–3288.

25. Hulsey-Vincent, H.J., E.A. Cameron, C.L. Dahlberg, and D.F. Galati, 2024 Spectral scanning and fluorescence lifetime imaging microscopy (FLIM) enable separation and characterization of C. elegans autofluorescence in the cuticle and gut. Biol Open 13 (12).

26. Klass, M.R., 1977 Aging in the nematode Caenorhabditis elegans: major biological and environmental factors influencing life span. Mech Ageing Dev 6 (6):413–429.

27. Kostich, M., A. Fire, and D.M. Fambrough, 2000 Identification and molecular-genetic characterization of a LAMP/CD68-like protein from Caenorhabditis elegans. J Cell Sci 113 (Pt 14):2595–2606.

28. Lam, A.B.Q., and J.M. Webster, 1971 Morphology and Biology of Panagrolaimus-Tipulae N Sp (Panagrolaimidae) and Rhabditis-(Rhabditella)-Tipulae N Sp (Rhabditidae), from Leatherjacket Larvae, Tipula-Paludosa (Diptera-Tipulidae). Nematologica 17 (2):201–212.

29. Laufer, J.S., P. Bazzicalupo, and W.B. Wood, 1980 Segregation of developmental potential in early embryos of Caenorhabditis elegans. Cell 19 (3):569–577.

30. Le, H.H., C.J. Wrobel, S.M. Cohen, J. Yu, H. Park et al., 2020 Modular metabolite assembly in Caenorhabditis elegans depends on carboxylesterases and formation of lysosome-related organelles. Elife 9.

31. Lee, H.J., W. Zhang, D. Zhang, Y. Yang, B. Liu et al., 2015 Assessing cholesterol storage in live cells and C. elegans by stimulated Raman scattering imaging of phenyl-Diyne cholesterol. Sci Rep 5:7930.

32. Luke, C.J., S.C. Pak, Y.S. Askew, T.L. Naviglia, D.J. Askew et al., 2007 An intracellular serpin regulates necrosis by inhibiting the induction and sequelae of lysosomal injury. Cell 130 (6):1108–1119.

33. Maupas, E., 1900 Modes et formes de reproduction des nematodes. Arch. Zool. Exp. Gen. 8:463–624.

34. Mendoza, A.D., N. Dietrich, C.H. Tan, D. Herrera, J. Kasiah et al., 2024 Lysosome-related organelles contain an expansion compartment that mediates delivery of zinc transporters to promote homeostasis. Proc Natl Acad Sci U S A 121 (7):e2307143121.

35. Morris, C., O.K. Foster, S. Handa, K. Peloza, L. Voss et al., 2018 Function and regulation of the Caenorhabditis elegans Rab32 family member GLO-1 in lysosome-related organelle biogenesis. PLoS Genet 14 (11):e1007772.

36. Nemys eds., 2025 Nemys: World Database of Nematodes, https://nemys.ugent.be.

37. Pincus, Z., T.C. Mazer, and F.J. Slack, 2016 Autofluorescence as a measure of senescence in C. elegans: look to red, not blue or green. Aging (Albany NY) 8 (5):889–898.

38. Rodak, N.Y., C.H. Tan, and P.W. Sternberg, 2024 An improved solid medium-based culturing method for Steinernema hermaphroditum. MicroPubl Biol 2024.

39. Roh, H.C., S. Collier, J. Guthrie, J.D. Robertson, and K. Kornfeld, 2012 Lysosome-related organelles in intestinal cells are a zinc storage site in C. elegans. Cell Metab 15 (1):88–99.

40. Schindelin, J., I. Arganda-Carreras, E. Frise, V. Kaynig, M. Longair et al., 2012 Fiji: an open-source platform for biological-image analysis. Nat Methods 9 (7):676–682.

41. Schroeder, L.K., S. Kremer, M.J. Kramer, E. Currie, E. Kwan et al., 2007 Function of the Caenorhabditis elegans ABC transporter PGP-2 in the biogenesis of a lysosome-related fat storage organelle. Mol Biol Cell 18 (3):995–1008.

42. Schroeder, N.E., and K.M. Flatt, 2014 In vivo imaging of Dauer-specific neuronal remodeling in C. elegans. J Vis Exp (91):e51834.

43. Schwartz, H.T., C.H. Tan, J. Peraza, K.L.T. Raymundo, and P.W. Sternberg, 2024 Molecular identification of a peroxidase gene controlling body size in the entomopathogenic nematode Steinernema hermaphroditum. Genetics 226 (2).

44. Schwarz, E.M., A. Baniya, J.K. Heppert, H.T. Schwartz, C.-H. Tan et al., 2025 Genomes of the entomopathogenic nematode Steinernema hermaphroditum and its associated bacteria. bioRxiv:2025.2001.2009.632278.

45. Sommer, R.J., L.K. Carta, S.Y. Kim, and P.W. Sternberg, 1996 Morphological, genetic and molecular description of Pristionchus pacificus sp n (Nematoda: Neodiplogastridae). Fundamental and Applied Nematology 19 (6):511–521.

46. Sommer, R.J., and P.W. Sternberg, 1996 Apoptosis and change of competence limit the size of the vulva equivalence group in Pristionchus pacificus: a genetic analysis. Curr Biol 6 (1):52–59.

47. Sternberg, P.W., and H.R. Horvitz, 1981 Gonadal cell lineages of the nematode Panagrellus redivivus and implications for evolution by the modification of cell lineage. Dev Biol 88 (1):147–166.

48. Sternberg, P.W., K. Van Auken, Q. Wang, A. Wright, K. Yook et al., 2024 WormBase 2024: status and transitioning to Alliance infrastructure. Genetics 227 (1).

49. Stiernagle, T., 2006 Maintenance of C. elegans. WormBook:1–11.

50. Stock, S.P., C.T. Griffin, and R. Chaerani, 2004 Morphological and molecular characterisation of Steinernema hermaphroditum n. sp (Nematoda: Steinernematidae), an entomopathogenic nematode from Indonesia, and its phylogenetic relationships with other members of the genus. Nematology 6:401–412.

51. Sun, F., Z. Zhao, M.M. Willoughby, S. Shen, Y. Zhou et al., 2022 HRG-9 homologues regulate haem trafficking from haem-enriched compartments. Nature 610 (7933):768–774.

52. Tan, C.H., H. Park, and P.W. Sternberg, 2022 Loss of famh-136/ FAM136A results in minor locomotion and behavioral changes in Caenorhabditis elegans. MicroPubl Biol 2022.

53. Tan, C.H., T.Y. Wang, H. Park, B. Lomenick, T.F. Chou et al., 2024 Single-tissue proteomics in Caenorhabditis elegans reveals proteins resident in intestinal lysosome-related organelles. Proc Natl Acad Sci U S A 121 (25):e2322588121.

54. Teramoto, T., and K. Iwasaki, 2006 Intestinal calcium waves coordinate a behavioral motor program in C. elegans. Cell Calcium 40 (3):319–327.

55. Thomas, J.H., 1990 Genetic analysis of defecation in Caenorhabditis elegans. Genetics 124 (4):855–872.

56. Thomas, L.J., and H. Quastler, 1950 Preliminary report of X-ray effects on the nematode Rhabditis strongyloides. Science 112 (2909):356–357.

57. Wang, H., K. Girskis, T. Janssen, J.P. Chan, K. Dasgupta et al., 2013 Neuropeptide secreted from a pacemaker activates neurons to control a rhythmic behavior. Curr Biol 23 (9):746–754.

58. Ward, A., J. Liu, Z. Feng, and X.Z. Xu, 2008 Light-sensitive neurons and channels mediate phototaxis in C. elegans. Nat Neurosci 11 (8):916–922.

59. Winter, C., 1992 The yolk polypeptides of a free-living rhabditid nematode. Comparative Biochemistry and Physiology. B, Comparative Biochemistry 103 (1):189–196.

60. Witte, H., E. Moreno, C. Rodelsperger, J. Kim, J.S. Kim et al., 2015 Gene inactivation using the CRISPR/Cas9 system in the nematode Pristionchus pacificus. Dev Genes Evol 225 (1):55–62.

61. Wong, W.R., K.I. Brugman, S. Maher, J.Y. Oh, K. Howe et al., 2019 Autism-associated missense genetic variants impact locomotion and neurodevelopment in Caenorhabditis elegans. Hum Mol Genet 28 (13):2271–2281.

